# IndivSTATIS: A multivariate approach to analyze brain network configurations with individualized parcellation

**DOI:** 10.64898/2025.12.19.695601

**Authors:** Ju-Chi Yu, Micaela Y. Chan, Liang Han, Erin Dickie, Hervé Abdi

**Affiliations:** Krembil Centre for Neuroinformatics, Campbell Family Mental Health Research Institute, Centre for Addiction and Mental Health, Toronto, ON, M5T 1R7, Canada; School of Behavior and Brain Sciences, The University of Texas at Dallas, Richardson, TX, 75080, USA; Center for Vital Longevity, The University of Texas at Dallas, Dallas, TX, 75235, USA; Department of Psychiatry, University of Toronto, Toronto, ON, M5S 1A1, Canada

**Keywords:** Multivariate analysis, Neuroimaging, Brain network configuration, Individualized parcellation, Functional brain connectivity

## Abstract

A critical step in the analysis of large-scale functional brain networks in neuroimaging is parcellation, which defines the nodes of a brain network. Group or atlas-based parcellation schemes use a shared common space, ensuring that each individual has the same number of brain parcels, which facilitates standard analytic approaches. However, studies reveal individual differences in the boundaries of brain areas. Extracting signals using atlas-based schemes can result in varying levels of blurring of signals across homogeneous areas within a specific individual’s brain. Individualized parcellation schemes can be obtained when sufficient data are available; however, these approaches introduce a significant analytical challenge: the number of parcels and networks differ across individuals. Here, we introduce IndivSTATIS, a new multivariate method based on the STATIS framework, designed to integrate individualized parcellation schemes while maintaining comparability across participants in a shared component space. The resulting network/node component scores can be used to predict individual differences measures (e.g., age, behavior). By allowing individualized parcellations to be compared within a common component space, IndivSTATIS provides a solution for incorporating individual network variability into larger studies, with potential to improve the sensitivity and interpretability of functional brain markers across both basic neuroscience and clinical applications.

## 1. Introduction

The brain operates as a large-scale network that performs information processing at multiple levels, from neurons, circuits, columns, areas, to large-scale functional systems that involve multiple, potentially distal, areas^1^. The prevalence of non-invasive human brain imaging, especially with functional magnetic resonance imaging (fMRI), has made it possible to investigate task-induced patterns of activation^2,3^ and the organization of large-scale functional networks^4,5^. Individualized brain parcellation has been proposed to define brain regions based on each individual’s own brain activity^6–8^, advancing the precision of human brain mapping. However, the lack of analytical methods to analyze individualized parcels has limited such efforts.

The functional organization of the human brain network is often examined using resting-state correlation^9^, where the covariation of blood-oxygen-level-dependent (BOLD) fMRI activity is used to infer functional relatedness between brain areas^10,11^. At the resolution of fMRI, brain areas can be identified using anatomically based parcellation (e.g., the AAL atlas)^12^ or functionally based parcellation (e.g., Gordon 333^13^ and Schaefer^14^ atlases). Functional-based parcellation typically uses clustering or boundary-detection algorithms, such as the watershed algorithm^15^, to detect functionally distinct areas (see **Fig. 1A** for an example of group parcellation from Gordon et al.^13^, which is also used in this work as the group atlas). These approaches often produce a parcellation scheme within a shared atlas space, to which individual brains are warped to the shared space (e.g., MNI152 in volumetric space; fs_LR in surface space). Using an atlas-based parcellation scheme, researchers often extract mean signals from each parcel and generate cross-correlation matrices that summarize the functional relationships between brain parcels. When using this approach, each individual’s brain uses the same parcellation atlas and has the same number of brain parcels or network nodes.

**Fig 1.**
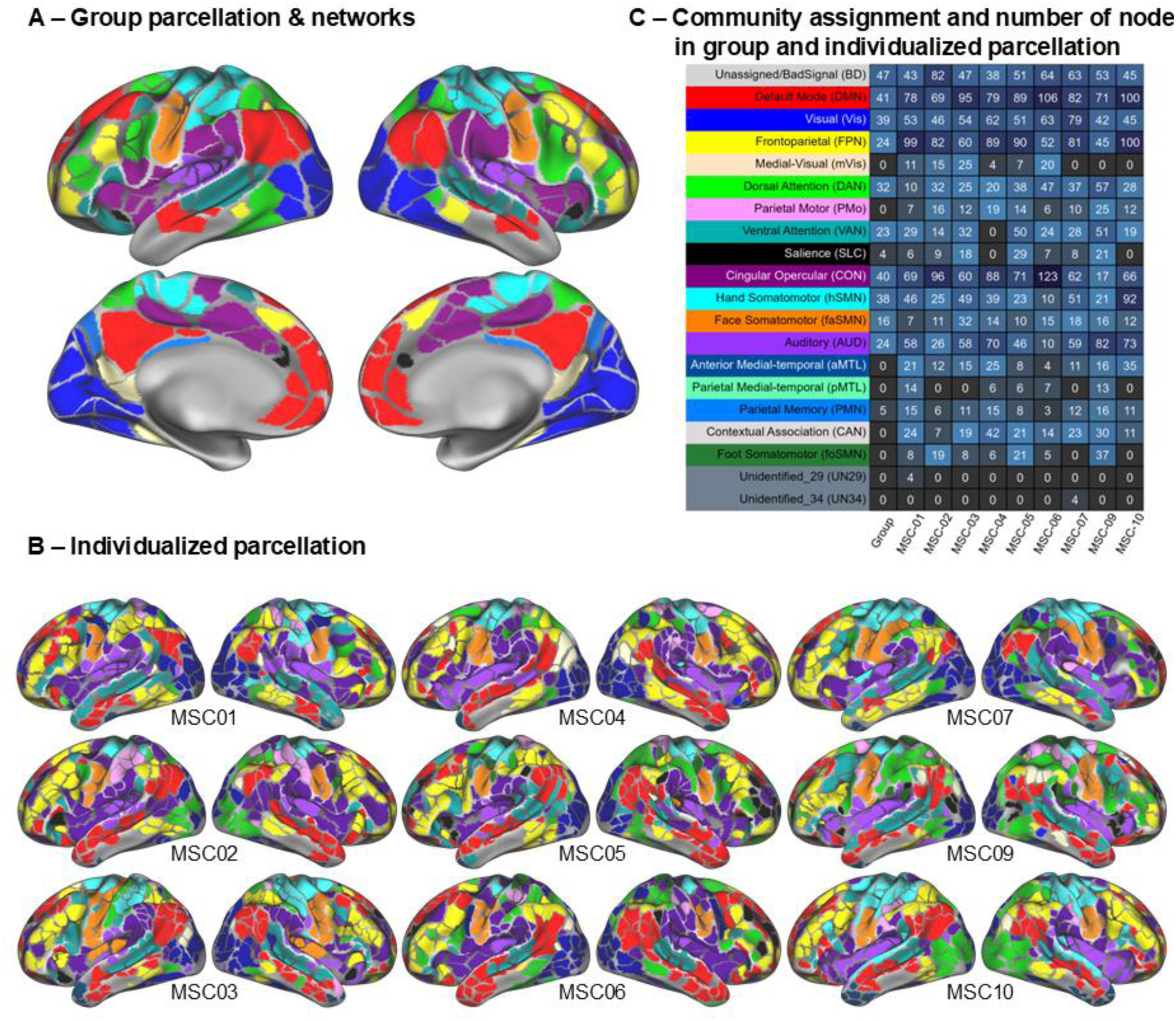
Individual-specific parcellations and networks differ not only across individuals but also from the group atlas. **(A)** Group parcellation from Gordon et al.^13^, generated using data from 120 young adults with boundary detection and the watershed algorithm. **(B)** Individualized parcellation using boundary detection mapping and the watershed algorithm for each participant. **(C)** Table showing the functional networks present for individual-specific parcellation per participant.

Studies have revealed individual differences in brain parcellation^7,8^ (see **Fig. 1B** for individualized parcellation from the Midnight Scanning Club (MSC) data presented in Gordon et al.^7^, which is used as the sample dataset in this work). Because of individual variation in brain area boundaries, extracting signals using atlas-based parcellations results in varying degrees of signal blurring between adjacent brain parcels. While using smaller “disks” or erosion away from a parcel border could mitigate some of these blurring, this approach still does not guarantee that the atlas-generated parcel is capturing signals from a homogeneous area within an individual’s brain. With sufficient data from a single individual—typically 20 minutes or more of resting-state fMRI after motion scrubbing—reliable functional network estimates can be obtained, consistent with prior work on network reliability^7,16^, and the broader finding showed that more resting-state functional connectivity (RSFC) data yields higher reliability^17^. Under these conditions, it is feasible to derive parcellation schemes that capture meaningful individual differences (e.g., ^6–8,18^). Despite these advances, many existing approaches for brain network analysis still face challenges in integrating individualized parcellations, making cross-subject comparisons difficult.

Common ways to compare brain networks across participants include comparing individual difference measures, such as age or clinical diagnoses^19–21^, network-based measures, such as modularity or segregation^22,23^, and node-level measures, such as degree or participation coefficient^24^. Unsupervised multivariate methods such as metric multidimensional scaling (MDS)^25^ and CovSTATIS^26,27^ (a three-way MDS; *Structuration des Tableaux à Trois Indices de la Statistique* [STATIS] for covariance matrices) can be applied to functional connectivity matrices derived from pair-wise correlation between BOLD signals from different parcels during an MRI scan and offer advantages in understanding functional network configurations (see examples using MDS in ^28,29^ and examples using CovSTATIS in ^30–32^). Other existing methods to analyze functional connectivity matrices across participants include approaches such as principal gradient analysis^33^, hyperalignment^34,35^, and Multi-session Hierarchical Bayesian model^36^.

Among these multivariate methods, MDS and CovSTATIS are two unsupervised methods that directly capture the variance in the connectivity patterns. MDS performs eigenvalue decomposition on a single connectivity matrix and extracts orthogonal components that explain the largest variance across the connectivity profiles of brain regions. With extracted components, one can visualize and quantify the configuration of brain parcels/networks based on the similarity between their activation over time. Such a configuration is often illustrated by scatter plots with axes representing the components and points representing the brain parcels.

CovSTATIS extends MDS to jointly analyze multiple connectivity matrices by extracting orthogonal components from the optimal linear combination of all matrices and projecting individuals’ connectivity patterns onto the same space to examine individual-level brain configurations. Among the more advanced analytical approaches, principal gradient analysis integrates geodesic brain features with functional connectivity and aligns each individual’s component space with a Procrustes rotation to extract the brain configuration across participants but does not allow the numbers of parcels to differ^33^; hyperalignment, instead of analyzing parcel-level information, extracts components from the voxel/vertex space, which is shared across participants, but is computationally heavy^34,35,37^; and Multi-session Hierarchical Bayesian model uses group atlases as a prior and allows the parcels to differ in shape, sizes, and the exact locations across participants while forcing the number of these extracted parcels to be the same^36^. However, all these aforementioned methods either (1) transform the participants who may have distinct parcellations into a shared parcellation/topographic scheme and require the number of parcels to be the same across participants or (2) analyze the participants with individualized parcellations separately (e.g., MDS), in which the resulting component spaces cannot be quantitatively compared.

To address these limitations, we developed ***IndivSTATIS***, a new multivariate method that integrates individualized parcellation while maintaining comparability across participants in the shared component space. IndivSTATIS is based upon applying the STATIS^26,38^ method to a data matrix generated by the cross-correlation of a data matrix generated with individualized parcellation with another data matrix created using atlas-based parcellation. STATIS stands for “*Structuration des Tableaux à Trois Indices de la Statistique*” (or, ‘structuring three-way statistical tables’ in English); it is also known as ‘*Analyse Conjointe de Tableaux (ACT)*’ (i.e., joint analysis of tables) or ‘*Procrustes matching by congruence coefficients*’^39^. The goals of STATIS and related methods are to (1) examine the similarity between different datasets (i.e., connectivity matrices in this case and in CovSTATIS), (2) to create an optimal weighted average—called the *compromise*—across all datasets that best represent the general pattern, and (3) to project each different dataset back into the space to allow comparisons by quantifying their discrepancies. IndivSTATIS adopts the same goals with specific aims to analyze connectivity patterns from individualized parcellation. Here, we first describe how IndivSTATIS can be used to examine brain network differences across participants with individualized parcellation. Next, we compare its unique effectiveness with existing unsupervised multivariate methods such as MDS and CovSTATIS.

The goal of the current work is to provide a framework that explicitly accommodates individualized parcellations while yielding a component space that is quantitatively comparable across individuals. Whereas other methods focus on improving alignment across individuals (e.g., hyperalignment, connectivity hyperalignment, hierarchical Bayesian models), the aim here is to preserve individualized parcellation structure and operate directly on individual-specific functional organization. Such flexibility in parcel/network numbers is especially important for improving the validity of research on populations with expected structural and functional differences, such as developmental or aging studies and clinical studies on participants with brain lesions or mental disorders.

## 2. Results

To compare and integrate the set of individual parcellations, ***IndivSTATIS*** combines group and individualized parcellations to create a common component space—known as a compromise—that places each participant’s individualized parcels and networks into a common frame of reference, enabling direct quantification and comparison across individuals. To illustrate IndivSTATIS (see **Fig. 2**), we use resting-state fMRI data from MSC (OpenNeuro: ds000224).

**Fig 2.**
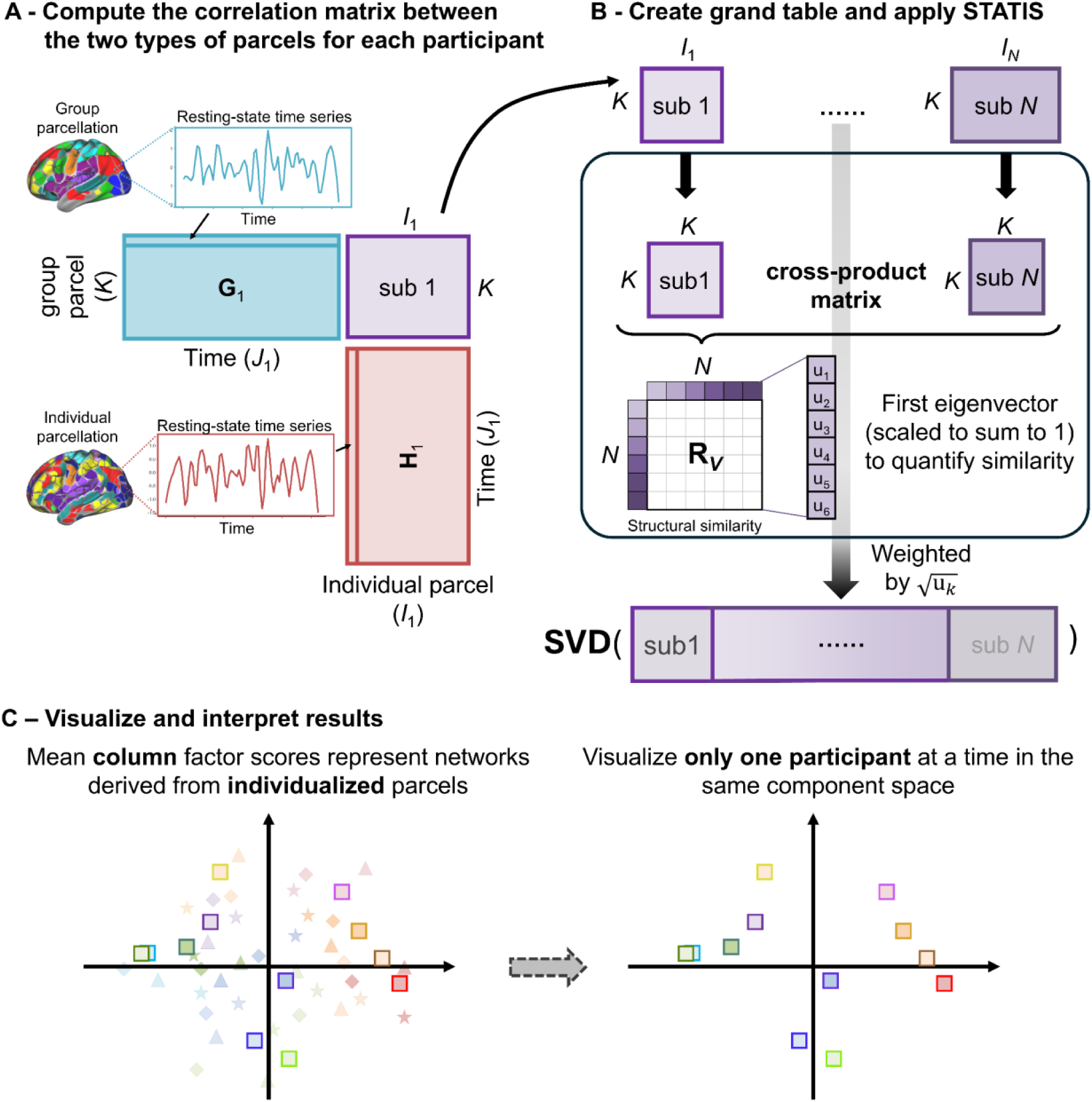
Illustration of IndivSTATIS. **(A)** We first extracted rsfMRI activation from both group and individualized parcellation; this process gives two matrices for each participant *n*: an *I_n_* by *J_n_* matrix **G***_n_* and a *K_n_* by *J_n_* matrix **H***_n_*. The cross-correlation of **G***_n_* by **H***_n_* gives an *I_n_* by *K_n_* participant block matrix, a matrix that describes the covariation between the group and the individualized parcels. **(B)** Next, IndivSTATIS normalizes each participant matrix by 1) their first eigenvalue (to normalize the variance across matrices) and 2) their structural similarity to the common pattern (so that the components are not driven by an outlier pattern). Finally, IndivSTATIS used the singular value decomposition (SVD) to extract the principal components that best characterize the variance across the individualized parcels (i.e., the column factor scores; left panel of **[C]**) according to their connectivity patterns, illustrating the functional network configuration. We can also show the subset of column factor scores of one individual to visualize the dissimilarity between their parcels (right panel of **[C]**).

IndivSTATIS (see **Fig. 2**) computes for each participant the correlations between two sets of time series: one set extracted from *K* parcels of a group parcellation (e.g., Gordon 333 parcels) and the other set extracted from *I_n_* parcels of individualized parcellation of Participant *n*. These *K × I_n_* correlation matrices, denoted by **X***_n_*, capture the similarity between each participant’s individualized parcels and their activations extracted using a common-group parcellation atlas. All these **X***_n_* matrices are then analyzed with STATIS^26^. With STATIS, each **X***_n_* is first normalized to have its first eigenvalue equal to 1 (an operation called MFA-normalization^40,41^) so that each participant contributes an equivalent amount of variance to the analysis. Next, these normalized matrices are weighted according to their similarity—measured by the *R_V_* coefficient^42^—to all the other **X***_n_* matrices. This step weights participants with a rarer pattern less and those with a more common pattern more, so that the derived compromise gives the best representation of the group. Finally, all weighted **X***_n_* are concatenated by their columns to build the grand table **X**, which is analyzed with the singular value decomposition (SVD)—an operation that expresses the **X** matrix as:

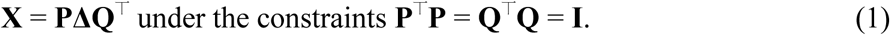

It is worth noting that the columns, representing the individualized parcels of all participants, are kept in the same scale and not normalized, such that the variation and the difference between zero and non-zero edges are preserved. From STATIS, the column factor scores are computed as *α_n_***QΔ** (where *α_n_* is the weight derived from the previous step for **X***_n_*). These column factor scores represent the individualized parcels of each participant on the compromise. See the **Methods** section for more details.

To compare across different group atlases, we show results using the Gordon 333 parcels^13^ and reanalyze them using two other atlases: the Chan 441 parcels^22^ and the Schaefer 400 parcels^14^ (results shown in the **Supplemental Information** [**SI**]). Additionally, we compare these results with IndivSTATIS using vertices or common networks rather than a predetermined group atlas. To further illustrate the utility of this method, we compared its results to the outcome of two existing methods: 1) an MDS analysis of each participant’s correlation matrix based on their individualized parcellation and 2) using CovSTATIS—a three-way MDS—of all correlation matrices with matching group parcels^26^.

### 2.1. IndivSTATIS component space

The column factor scores of all individualized parcels are shown in **Fig. 3** as a scatter plot with axes corresponding to (orthogonal) components. After averaging across networks within each participant, we showed the mean column factor scores of each individualized network of each participant in **Fig. 3**. Participant 1’s network factor scores are highlighted, along with those from other participants. As the participants are all in the same space, we can compare the network configuration across participants with their individualized parcellation/networks. It is worth noting that the IndivSTATIS results presented in the main text kept both positive and negative edges for further analysis. For comparison, we included IndivSTATIS results without negative edges in **Fig. S1** in the **SI**.

**Fig 3.**
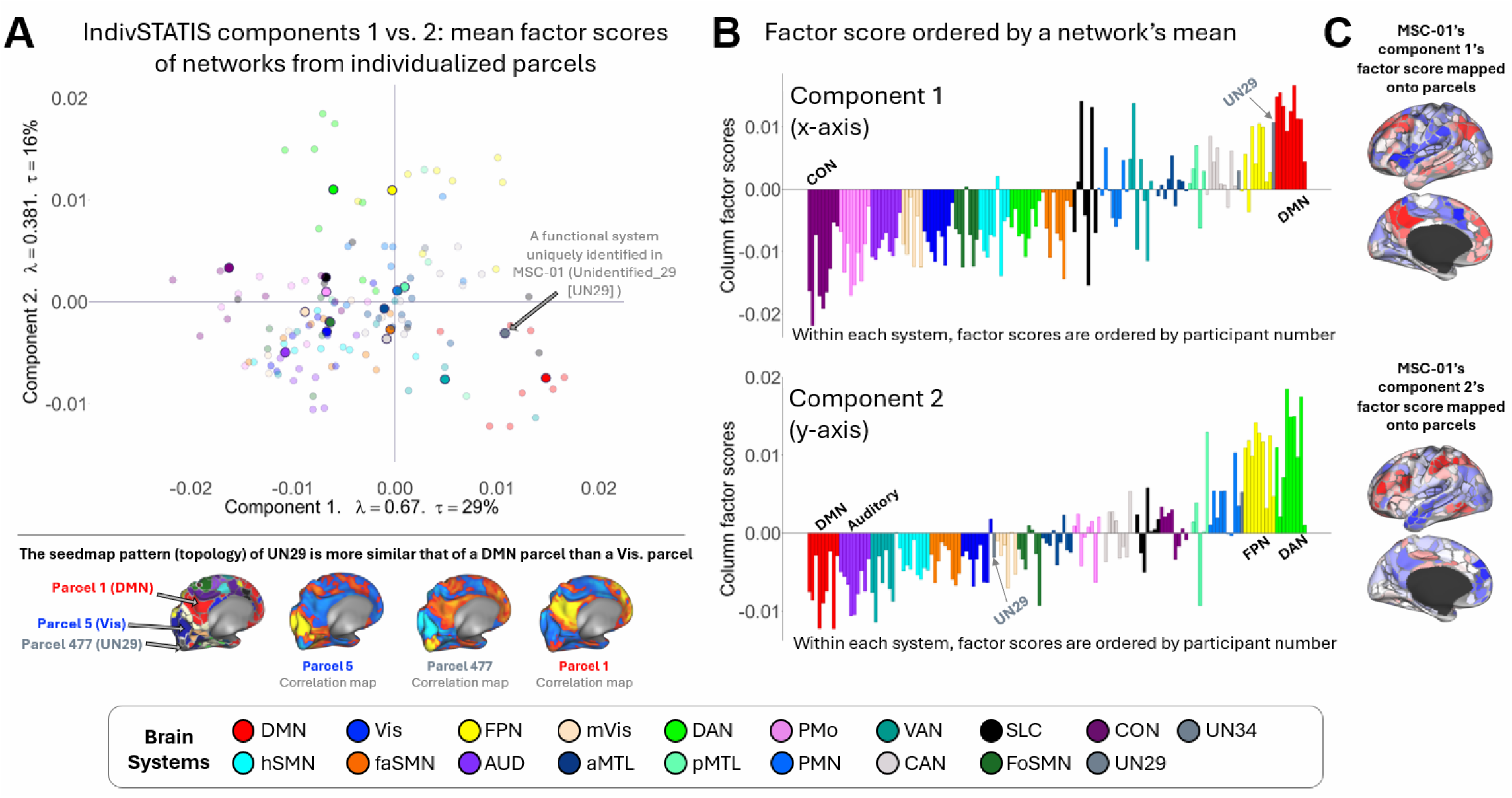
IndivSTATIS Components 1 and 2 reveal individual variability in network patterns. **(A)** Participants’ mean column factor score for each functional network is plotted along IndivSTATIS Components 1 and 2. To single out one participant, MSC-01’s functional network’s mean column factor scores are plotted as large and solid circles; all the other participants are plotted as smaller and transparent circles. Component 1 is characterized by the *opposition*, indicating the anticorrelation, of two association networks: cingulo-opercular (CON; dark purple) and default mode network (DMN; red; see Fig. 1 for the topographic location of CON and DMN across participants). One of the key features of this method is its ability to visualize functional networks unique to an individual. Unidentified 29 (UN29 in grey) is highlighted by an arrow on the right side of the biplot. This is a network uniquely identified in MSC-01. It is situated in the occipital pole (see brain resting-state correlation maps below, biplot highlighting three parcels: DMN, Vis, UN29), but exhibits a cortical correlation map resembling DMN instead of Vis or mVis (correlation brain map that belongs to a UN29 parcel is more similar to a DMN parcel’s correlation map than a Vis parcel’s correlation map). **(B)** The barplots (top: Component 1; bottom: Component 2) sort each participant’s mean factor score by (i) network and (ii) participant number. In Component 1, CON [purple] is most negative across all participants, so it is grouped together on the left-most side of the barplot, and MSC-01’s CON mean factor score is the first purple bar. Again, here UN29 is situated close to DMN. **(C)** Each parcel’s factor score is visualized on the cortical surface.

Column factor scores from IndivSTATIS (**Fig. 3A**) showed that the network configuration in component space is similar across participants. Together, the top six components explain 63% of the variance. Component 1 (proportion of variance explained, *τ* = 25%), shown in the horizontal axis of **Fig. 3A** and the top panels of **Fig. 3B-3C**, is characterized by the opposition of the default mode network (DMN) and cingulo-opercular network (CON). Component 2 (*τ* = 16%), shown in the vertical axis of **Fig. 3A** and the bottom panels of **Fig. 3B-3C**, differentiates the dorsal attention (DAN) and frontoparietal networks (FPN) from all other networks.

The association networks — such as the cingulo-opercular (CON), frontoparietal (FPN), ventral attention (VAN), contextual association (CAN), and default mode networks (DMN) — create more distinct clusters on the factor map, whereas the sensorimotor networks — such as auditory (Aud), visual (Vis), hand and face somatomotor networks (hSMN and fSMN) — overlap with each other. These networks show less interindividual variability than memory-related networks, such as parietal memory (PMN), anterior and posterior medial temporal (aMTL and pMTL), and salience networks (SLC).

In addition to the comparison between the general functional brain configuration, IndivSTATIS can identify brain parcels that were not categorized as known functional networks. For example, the specific parcel from Participant 1 is labeled ‘Undefined 29’ (parcel 447; shown in **Fig. 3A**), because the algorithm did not categorize it into a known network. This parcel is topographically located in the occipital lobe, which is typically part of the visual network, but is functionally more similar to DMN (see the Parcel 477 correlation map in **Fig. 3A**; for reference, separate correlation maps from Parcel 1, a DMN parcel, and Parcel 5, a Vis parcel, are shown in the bottom corners of the figure). Its topographical location in the occipital lobe and its moderate connectivity mostly to vertices in close topographic (physical) proximity to the DMN result in the algorithm placing it in a distinct community. In analyses that use only group-parcellation/communities, including this parcel as part of the visual network will create a noisier visual network connectivity pattern.

### 2.3. Interpretation

#### 2.3.1. Interpreting factor scores

Conceptually, the *compromise* optimally aligns the group parcels across participants and projects the individualized parcels of all participants onto the same component space—an operation providing a metric to quantify and compare these individualized parcels or networks.

The extracted components of the compromise are defined by the opposition of the networks from the two extreme ends. These oppositions of networks illustrate the most characteristic aspects of the functional brain network’s configuration. The parcels are represented on these components as factor scores, whose values quantify how functionally connected a parcel is to the two opposing network sets in a given dimension. The functional similarity between any two parcels can be quantified by the cosine of the angle formed by these two parcels in the compromise with respect to the origin (i.e., cos 0° = 1, cos 90° = 0, cos 180° = -1). In addition, we can take the mean factor score of each network to illustrate the network-level configuration of each participant.

Mean factor scores can also be computed for each parcel or each network across groups of participants to examine group effects in the brain’s functional organization. For example, mean network factor scores can be correlated with cognitive measures to examine how network configuration relates to cognition, or can be compared across groups to reveal differences from distinct populations, such as sex, healthy vs. clinical groups, age, etc.

Other metrics can be extracted from this component space, such as the area covered by parcels of a given network in this component space. The size of this area reflects the network’s homogeneity, with a smaller area indicating a more homogeneous network. Such a measure of homogeneity can be computed for each participant or across a group of participants to further examine individual differences in brain configurations. For example, a correlation between age and the network homogeneity values of the DMN can demonstrate the aging effect of DMN in an aging sample. Overall, with IndivSTATIS, participants are allowed to have different networks, while each of them may also include different numbers of parcels.

##### 2.3.1.1 Example of interpreting factor scores

In this section, we show an example of how results from IndivSTATIS can be analyzed to examine their relationships to other measures. We used performance on the Picture Vocabulary Task of the NIH Toolbox^43^ and age (in years) from the MSC datasets to demonstrate how we can examine behavioral and demographic relationships with brain network configuration. We performed Pearson’s correlations between different IndivSTAITS outputs to either the behavioral performance or age. Due to the small sample size (*N* = 9), we will not interpret the significance level but focus on the effect size by considering an *r* > .50 indicative of a meaningful association. It is worth noting that we selected this specific task from the MSC dataset because it showed the clearest and most interpretable effect, solely to show how to interpret results from IndivSTATIS. This task was not chosen based on theoretical considerations, and therefore, the interpretations presented below should not be considered scientific conclusions, especially given the small sample size.

In **Fig. 4A**, we computed the mean network factor scores for each participant’s available network and correlated them with Picture Vocabulary Task performance, with higher task scores indicating better performance in choosing the visual stimuli that matched the heard vocabulary. For easy interpretation, we first defined the two identified components. We called Component 1 the “*Attention component*”, as it was characterized by a gradient from a focused attention end of the axis (represented by CON)^44^ to the other end of medium-level (represented by FPN involved in task switching^45^) or free attention (represented by DMN)^46^. We called Component 2 the “*Attention Control component*”, as it was characterized by the opposition of FPN (involved in adaptive attention control), DAN (involved in top-down, goal-oriented attention control) versus DMN (involved in internal processing), and VAN (involved in bottom-up, stimulus-driven attention control)^47^. Results showed that better Picture Vocabulary Task performance is related to less differentiated DMN from CON and the auditory network (AUD) along the Attention component; correlations between the performance and the Attention Control component are weaker, with a mixed pattern.

**Fig 4.**
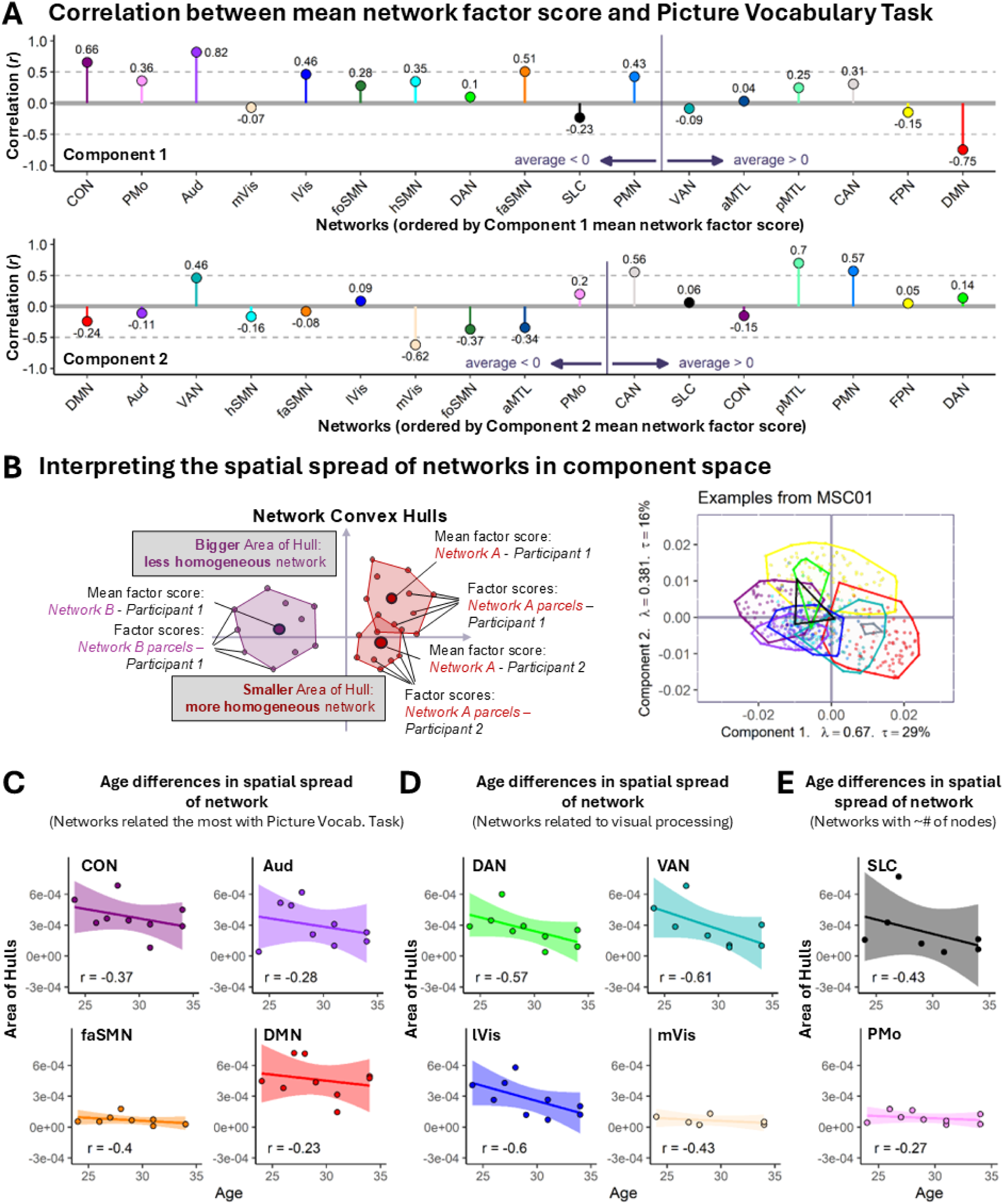
IndivSTATIS factor score and spatial spread (area of hull) is associated with behavior and age. **(A)** Correlation between each network’s mean factor score and performance on the Picture Vocabulary task (top: component 1; bottom: component 2). Networks along the x-axis are ordered by the networks’ mean factor scores within each component. **(B)** The spatial spread of the factor scores in component space could be encircled by the “area of hull”. A bigger area of hull is associated with a less homogeneous network (factor scores within a network are further apart), and a smaller area of hull is associated with a more homogeneous network. Correlation between age and each network’s area of hull (see panel B for interpretation) across MSC participants. Networks most strongly associated with the Picture Vocabulary task are grouped in **(C)**, and those linked to visual processing in panel **(D)**. To demonstrate that these associations are not simply driven by the number of nodes within a network, **(E)** compares the SLC and PMo networks, which have similar numbers of nodes but differ in their age–area-of-hull relationship.

We can also quantify the homogeneity of a given network by computing the area size of the convex hull spanned by all its parcels in the component space; as shown in **Fig. 4B**, a larger convex hull indicates a less homogeneous network. In **Fig. 4C-4E**, we showed three different correlation analyses examining the age effects (from 24 to 34 years old) of network homogeneity. **Fig. 4C** illustrates that the four networks with the strongest correlation to Picture Vocabulary Task performance also showed medium-level age effects, where participants in their 30s tend to have more homogeneous networks than those in their 20s. **Fig. 4D** illustrated that the four networks related to visual processing showed strong age effects, where participants in their 30s tended to have more homogeneous networks than those in their 20s. Finally, **Fig. 4E** showed the comparison between the salience network (SAL; having 6-25 parcels across participants) and the parietal motor networks (PMo; having 4-29 parcels across participants). These two networks have similar numbers of parcels, with SLC showing a strong age effect and PMo showing a medium-level age effect of network homogeneity, implying that similar sizes of individualized functional networks can lead to different levels of effect in network homogeneity.

#### 2.3.2. Comparison to other component-based methods

To characterize IndivSTATIS, we compared it to two other similar approaches: MDS and CovSTATIS. To properly compare MDS and CovSTATIS with the IndivSTATIS results, we performed MDS and CovSTATIS on functional connectivity matrices with Pearson’s correlation coefficients (*r*) without Fisher’s *Z*-transformation and double-centering; the results are shown in **Fig. 5** and **S2**. For simplicity, only the results from two participants, MSC-01 and MSC-02, are shown in **Fig. 5**, and results across all 9 participants from the full sample are included in **Fig. S2**. To examine how well IndivSTATIS captures the configuration at the individual level, we quantify the structural similarity (i.e., matrix similarity beyond rotation and scaling) with an *R_V_* score, a measure analogous to an *R*^2^ (for the dimensional structure similarity between matrices), between factor scores from IndivSTATIS and from the corresponding MDS.

**Fig 5.**
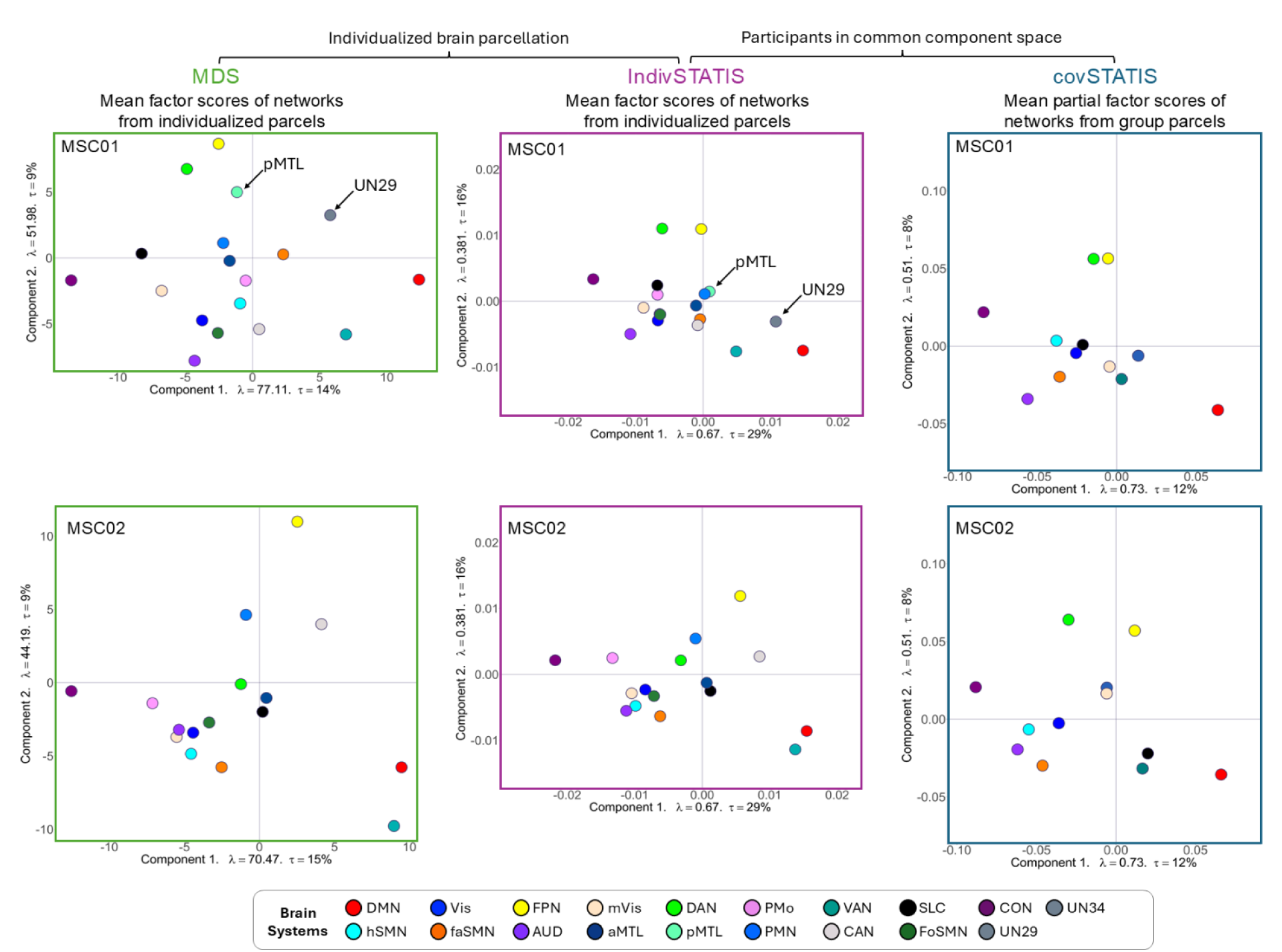
IndivSTATIS combines the advantages of MDS (displaying individualized networks) and CovSTATIS (plotting individuals in the same component space). Component biplots for two participants, MSC-01 (top) and MSC-02 (bottom), are presented to illustrate the differences in network configuration between these methods. This figure shows the results analyzing the data derived from the same fMRI scan using multidimensional scaling (MDS; left column), IndivSTATIS (middle column) with individualized networks, and covSTATIS (right column) with networks derived from a group atlas (i.e., Gordon et al.^13^). Each round dot illustrates the mean factor score of each network and is colored accordingly. Networks that are unique to MSC-01 (pMTL & UN29) only appear on MDS/IndivSTATIS (left and middle top row). MDS performs separate decompositions on participants’ connectivity matrices and derives different component spaces. For instance, the two left (green) scatter plots of different participants do not have the same component space (note the difference in axis), their components therefore have different eigenvalues and proportions of explained variance. MDS allows individualized parcels/networks but limits the comparisons as the spaces differ. For IndivSTATIS and CovSTATIS, the two scatter plots of different participants illustrate brain configurations in the same component space. The axis labels, therefore, show the same eigenvalues and proportions of variance explained within each method. However, CovSTATIS requires the same node set and cannot examine unique networks; therefore, the pMTL and UN29 networks that are not common across all participants are only available in the IndivSTATIS results (see arrows) but not covSTATIS.

In MDS (left column of **Fig. 5**), each participant was analyzed separately, yielding individual-specific component space (note different Components 1 and 2 axes between MSC-01 and MSC-02’s MDS space). Therefore, individualized networks (e.g., pMTL and UN29 in MSC-01) can exist, and each individual’s component space is shown in a separate plot. Individuals’ component space extracted from separate MDS precludes direct comparison of factor scores across participants. The IndivSTATIS results are visualized in **Fig. 5** and demonstrate close structural similarity between its factor scores and factor scores from individuals’ MDS (*R_V_* range = [.87, .94], MSC-01: *R_V_* = .87, MSC-02: *R_V_* = .87, see comparisons of other participants in **Fig. S2**).

In CovSTATIS (right column of **Fig. 5**), all 9 participants were analyzed together and contributed to the same component space, and we showed the mean network factor scores in two separate plots for MSC-01 and MSC-02 (note the same Components 1 and 2 axes for both participants). When plotted together, the cosine of two networks’ or parcels’ coordinates, even between individuals, can be used to describe the functional similarity/difference of those networks/ROIs. However, CovSTATIS requires the input network to have the same node and network assignments. Therefore, networks with distinct topographic/topological patterns, such as UN29, will not emerge in analysis utilizing CovSTATIS.

In IndivSTATIS (middle column of **Fig. 5**), while the component space is also defined by all the participants (as in CovSTATIS), the configurations of subnetworks that are distinct to specific individuals are also retained (as in MDS). In sum, IndivSTATIS provides a good consensus that captures both the common and the individualized patterns from precision brain mapping.

#### 2.3.3. Comparison across different group atlases or references

To examine whether our results are specific to the Gordon 333 atlas^13^, we explored other options for group atlases, Chan 411 atlas^22^ and Schaefer 400 parcels atlas^14^. We first used Chan atlas^22^—which derived uniform disks instead of parcels—to examine a group atlas which was built with a similar algorithm and parcel numbers to Gordon et al.^13^, but that have parcels of equal sizes. The results are shown in **Fig. S3A** and reveal close structural similarity between its factor scores and the factor scores from the individuals’ MDS (*R_V_* range = [.82, .93]) and from IndivSTATIS that used the Gordon 333 atlas^13^ (*R_V_* range = [.96, .99]). For an alternative group atlas that used a ^d^ifferent parcellation algorithm, IndivSTATIS results from the Schaefer 400 parcels atlas^14^ are shown in **Fig. S3B**. The results showed close structural similarity between its factor scores and factor scores from individuals’ MDS (*R_V_* range = [.87, .94]) and from IndivSTATIS that used the Gordon 333 atlas^13^ (*R_V_* range = [.84, .92]).

For alternative alignment references which are free from any existing group parcellation template, we examined two group alignments that have high and low spatial resolution, respectively: the vertices and the common networks. In the vertex approach, IndivSTATIS analyzes Pearson’s correlations between time series from each vertex and from the individualized parcels. The results (displayed in **Fig. S3C**) show that the brain configurations derived using vertices are also similar to the actual individual factor space extracted by both the MDS (*R_V_* range = [.85, .93]) and IndivSTATIS using the Gordon 333 atlas^13^ (*R_V_* range = [.96, .99]). In the common network approach, IndivSTATIS analyzed Pearson’s correlations between the mean time series from the 9 common networks across all participants and time series from the individualized parcels. The results (see **Fig. S3D**) show that the brain configurations derived using common networks are also similar to the actual individual MDS factor space (*R_V_* range = [.87, .95]) and factor space derived by IndivSTATIS using the Gordon 333 atlas^13^ (*R_V_* range = [.94, .99]). As the vertex approach is computationally heavy, if one wants to be completely free of group templates, common networks are a better option than vertices to achieve the same similarity to individuals’ MDS while aligning participants.

#### 2.3.4. Individual differences in comparison with the existing graph-theory-based network measures

To validate and understand IndivSTATIS, we compared IndivSTATIS to existing graph-theory-based metrics that have been traditionally used to characterize functional connectivity patterns. These metrics include participation coefficients (PC) and within-module *Z* scores (WMZ) for each brain area, as well as the segregation of each network. PCs ranging between 0 and 1 measure how connected each brain region is to another network compared to the network it is in; the higher the value, the more connected it is outside of rather than within its own network.

WMZs measure how connected each brain region is to other nodes within its network; the higher the value, the more connected the region is to other within-network regions. Finally, segregation describes how disconnected a network is from all other networks in the brain.

To help understand the space of IndivSTATIS, we computed PC and WMZ for each individual’s parcels and the segregation of each individual’s networks from connectivity matrices thresholded at 4%. Results showed that PC is quadratically related to the factor scores of Components 1 & 2 from IndivSTATIS (**Fig. 6A**), where the brain region with a higher absolute value is likely to have a lower PC (i.e., connected more within than outside of its own network). WMZ were also quadratically related to the factor scores, but for both Components 1 and 2 from IndivSTATIS (**Fig. 6B**), where the brain regions closer to the origin of the component space have lower WMZ scores (i.e., less connected to other regions within their own networks). Finally, the segregation of a given network is positively related to the average squared distances from this network to all other networks (of the same participant) in the component space (see **Fig. 6C**). These relationships are illustrated by the diagram in **Fig. 6D**. In sum, the component space not only visualizes but also depicts the brain configuration of individualized parcels, quantifying how these brain regions are functionally connected to one another.

**Fig 6.**
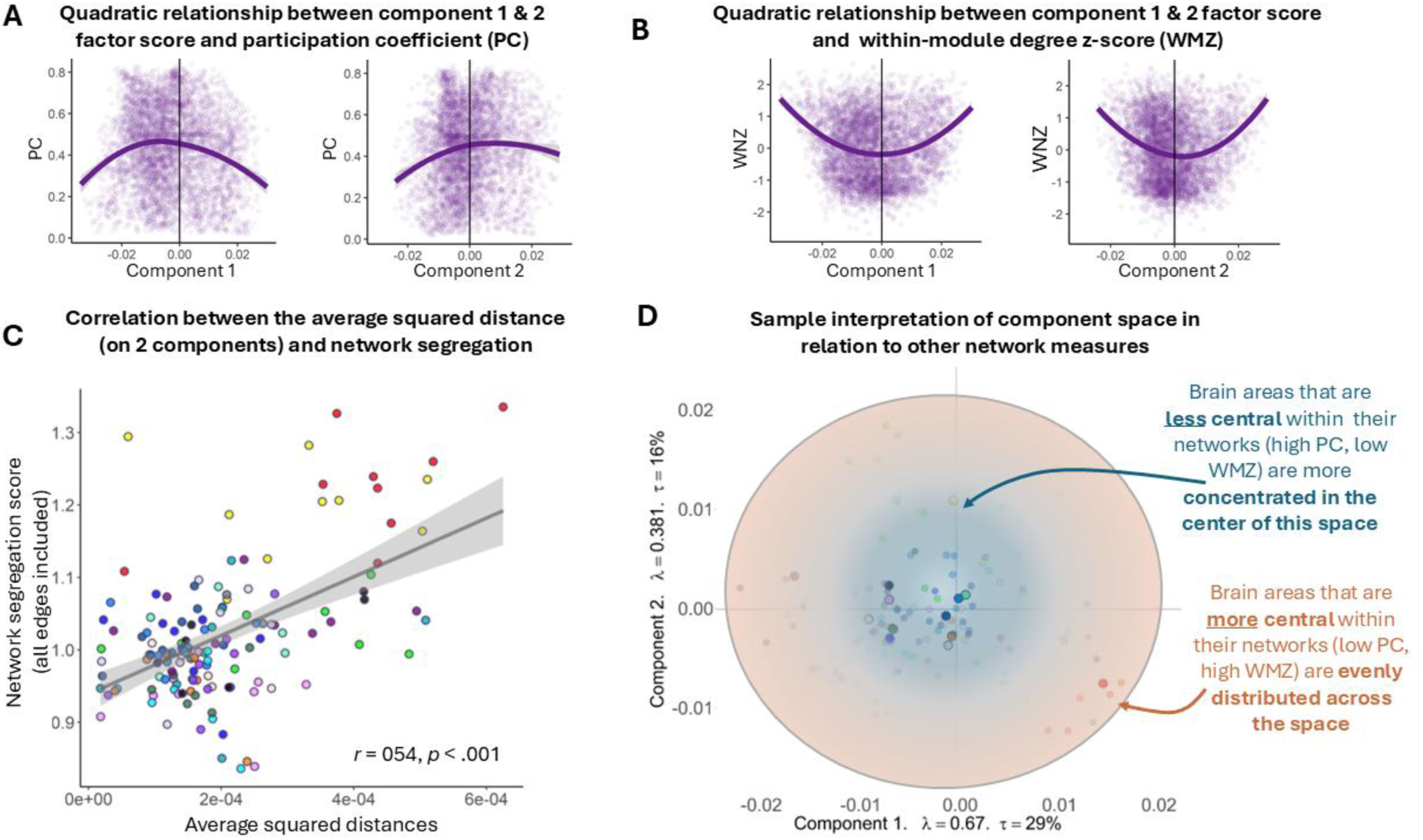
Relationship of IndivSTATIS Components to other network measures. Across all participants, each individualized brain area’s factor score is related to (**A**) participation coefficient and (**B**) within-module *Z*-scores. **(C)** Sum of squared distances of an individual’s sub-network (each dot is a sub-network for one MSC individual). Sub-networks with a high sum of distance in the IndivSTATIS component space have higher network segregation. **(D)** Diagram illustrating that the center of the component space (of Components 1 and 2) corresponds to brain areas that exhibit more out-of-network (less constrained) functional correlation with other brain areas, which likely correspond to brain areas that are less central to its subnetwork.

## 3. Discussion

We present a new statistical technique to compare brain networks derived from different individualized parcellation, where the number of network nodes differ across participants. This method uses group references (here, group-based parcellation) to align brain networks derived from individualized parcellations, where the number of network nodes varies across participants. Previously, using individualized parcellation typically restricted researchers to only compare summary measures of the whole network or subnetworks (e.g., computing network modularity or segregation). While multivariate techniques such as MDS provide a customized component space for each individual brain network, the actual components are not comparable across participants. Other multivariate methods, such as CovSTATIS, create a component space that is shared across participants, but necessitate the same number of network nodes (i.e., rows and columns of the matrix) for each participant.

To overcome these limitations, the proposed method—IndivSTATIS creates a shared component space by using both the individualized parcellations and a group-based parcellation as a common alignment reference—a procedure effectively aligning individualized networks into a common representational space. This procedure enables direct comparison across participants, while still preserving individual-specific variations in brain network topography and topology. Importantly, while we chose the Gordon parcellation, the exact choice of group-based parcellation does not significantly alter the results (see **Fig. S3A**). In the **SI**, we demonstrate that the group parcellation derived using alternative clustering algorithms (e.g., gradient-weighted Markov Random Field used to generate the Schaefer 400 node set^14^) or topographically uniform shapes (e.g., geodesic disks used by Chan et al.^22^) yield similar results. This pattern suggests that IndivSTATIS is robust to different parcellation strategies if the node sets capture sufficiently strong and reliable signals.

In addition, if one wants to be completely free of a group atlas, we further showed that the vertices from the fs_LR 32k surface and common networks shared across participants can also be a good alignment reference. However, these non-atlas based alignments should be implemented with caution, especially when imaging quality is low (e.g., scans less than 20 minutes), because time series from vertices may introduce more noise and be too unstable to serve as a reliable group alignment source, and the common networks, on the other extreme, could yield weaker results than using a group-based parcellation.

The IndivSTATIS approach is particularly useful for clinical and developmental research where individual variations in brain anatomy and functional network topology are central to these questions. IndivSTATIS enables meaningful comparison across participants while retaining individual-specific features, a feature making it possible to identify personalized markers of development or disease trajectories that would otherwise be obscured in methods requiring shared node definition. IndivSTATIS offers researchers a flexible framework for uncovering clinically or developmentally relevant individual differences in brain network topology, and will help shed light on deep phenotyping of heterogeneous clinical populations.

### 3.1. Comparison to other methods

The two major goals of IndivSTATIS are to (1) align participants who have different numbers of parcels/networks into a shared space, and (2) extract components that help understand brain network configuration within and across participants. In the present manuscript, we compared results from IndivSTATIS to results from MDS and CovSTATIS because these three methods have relatively similar approaches to decomposing data (i.e., linear-based decomposition using SVD). The results show that IndivSTATIS retains individual-specific network characteristics—as also shown in the MDS results—but also successfully brings all individuals into a common space.

Other methods and approaches exist to address one of the two issues that IndivSTATIS aims to solve. For example, another notable alignment method, such as *hyperalignment*, captures fine-grained functional correspondence across participants^34,37^. Application of hyperalignment to resting-state connectivity aligns participants based on connectivity profile instead of task-evoked responses^35^. However, hyperalignment requires consistent dimensionality (number of nodes or voxels). When applied to connectivity analysis with individualized parcellation, it requires vertex-wise connectivity, which typically requires large datasets for reliable alignment and is computationally demanding. By contrast, IndivSTATIS provides an efficient and interpretable solution for individualized parcellations, as long as reasonably a reliable group-based reference can be used to align individuals.

Other common approaches to extracting components from brain networks data is CovSTATIS (which we covered above), as well as gradient based analysis^33^, which have been used extensively in recent times to understand patterns of brain organization related to development and aging^48–50^, as well as various diseases and disorders^51–54^. However, most of these component extraction methods require consistent dimensions across participants. **Table 1** outlines some of the common methods/approaches used to address individual variations and/or utilize component spaces in analysis, highlighting why these various approaches are solving important but different issues.

**Table 1.** Summary and comparison with other alignment/component methods.

| <u>Method</u> | <u>What it does</u> | <u>What it doesn't do</u> |
| --- | --- | --- |
| **Goal: analyze data in component space** |  |  |
| **metric/Classical MDS<sup>25</sup>** | Creates a component space for each individual network of different sizes. | Does not align networks with different parcellations to the same space for analysis. |
| **CovSTATIS<sup>26,27</sup>** | Creates a shared component space across networks, but requires the same size matrix. | Does not accommodate matrices of different sizes. |
| **Gradient**<sup>33</sup> | Similar to CovSTATIS, it extracts components and handles multiple connectivity matrices (nonlinear), | Does not accommodate matrices of different sizes. |
| **Goal: align activity patterns into a common functional space based on functional response** |  |  |
| **Hyperalignment**<sup>34,37</sup> | Requires a reference task(e.g., movie-watching) with similar stimuli.<br>transforms activity to common space using high-dimensional *procrustes* transformation /ridge regression<br>can be used to create more reliable functional parcellation and network | Does not use *individualized parcellation*. This method addresses the ‘individualized’ issue by analyzing vertices and aligning them with Procrustes rotation. |
| **Connectivity Hyperalignment (CHA)**<sup>35</sup> | Instead of aligning evoked response, it aligns the functional network structure (vectorized connectivity pattern) | Does not use *individualized parcellation*. This method addresses the ‘individualized’ issue by analyzing vertices and aligning them with Procrustes rotation. |
| **Goal: improve parcellation** |  |  |
| **Hierarchical Bayesian Brain Parcellation Framework**<sup>18</sup> | Combines information from different datasets to achieve an improved population-based atlas.<br>Generates individual parcellation using fewer amount of data (e.g., 10 min of data) | Does not align different parcellations to the same space for analysis<br>Does not allow individualized networks |
| **Multi-session hierarchical Bayesian model (Areal MS-HBM)**<sup>36</sup> | Generates parcellation based on priors, with flexibility to adjust the number of parcellations based on participants, resulting in high-quality parcellation (higher homogeneity) while | Does not allow individualized networks<br>Imposes constraints on the number of individualized parcels |
|  | maintaining the same number of parcellations across participants |  |

### 3.2. Limitations

Despite its advantages, IndivSTATIS has certain limitations. While our findings suggest that IndivSTATIS is robust when using different group-based parcellations for alignment (see **Fig. S3** showing similar results using Schafer 400 and Chan 441 atlases), further research is needed to assess how template selection affects specific network-level inferences. Also, while IndivSTATIS is effective in aligning individuals to a shared space, it does not improve the quality of the individualized parcellation. Thus, any issues that impact the quality of the individualized parcellation will be propagated in this style of analysis, such as insufficient data quantity, types of scan sequences (multi-echo scans may reduce amount of data needed^55,56^), image artifacts, and other preprocessing choices. Additionally, IndivSTATIS only captures the linear relationships between the networks—a feature which is a strength in terms of interpretability (e.g., reconstructing data), however, future extensions could incorporate nonlinear relationships.

### 3.3. Conclusion

In conclusion, the present work advances brain network analysis by introducing and illustrating IndivSTATIS—a new statistical method that balances individualized precision and cross-participant comparability. By integrating individualized and atlas-based parcellation schemes into a common statistical framework, IndivSTATIS offers a robust and scalable solution for understanding network variability in large-scale neuroimaging datasets. Future work will explore the applicability of this method to longitudinal and intervention data (directly see how the distance between data in the biplot reflects experimental manipulation) and in larger datasets to explore individual differences.

## 4. Materials and Methods

### 4.1. Data Description

Data from the Midnight Scanning Club (MSC)^7^ were used in the present study. Here, we briefly summarize the data parameters and processing. Brain images were collected on a Siemens TRIO 3T MRI scanner from 12 sessions, on separate days. Anatomical images were collected on two separate days. A total of four T1-weighted images (TR = 2400 ms, TE = 3.74 ms, TI = 1000 ms, resolution = 0.8 × 0.8 × 0.8 mm^3^, flip angle = 8°, 224 slices) and four T2-weighted images (TR = 3200 ms, TE = 479 ms, resolution = 0.8 × 0.8 × 0.8 mm^3^, 224 slices) were obtained for each participant across two separate days. Resting-state functional MRI images were acquired across 10 sessions (1 session per day). In each session, 30 minutes of resting-state fMRI (gradient-echo EPI, TR = 2.2 s, TE = 27 ms, flip angle = 90°, resolution = 4 × 4 × 4 mm^3^, 36 slices) were collected while the participants kept their eyes open and fixed on a white fixation cross against a black background. One participant (MSC-08) with poorer quality data was excluded from the present analysis^7^.

### 4.2. Preprocessing

Mean anatomical images (T1- and T2-weighted) were derived by averaging structural images with the same modality. The mean field map was used to correct for distortion for each participant^7,57^. An individual’s native surface was generated using the following steps: (i) FreeSurfer v5.3 was used to generate anatomical surfaces from the participant’s average T1-weighted image, (ii) the fsaverage-registered surfaces from the FreeSurfer pipeline was registered and resampled to a hybrid atlas surfaces with 163,842 vertices (164k fs_LR)^58^ using landmark-based deformation, performed using Caret tools^59^, (iii) the 164k fs_LR surface data were down-sampled to a 32,492 vertices surfaces (fs_LR 32k). A one-step sampling was performed by combining the deformations from the participant’s native surfaces to the fs_LR 32k surfaces into a single deformation map.

The functional BOLD images were processed by performing basic fMRI preprocessing: (i) slice-timing correction, (ii) intensity normalization^60^, (iii) realignment, (iv) registration and resampled to the Talairach template^61^ in 3mm isotropic space.

Resting-state BOLD signal requires additional processing steps to reduce the impact of head motion on the data: (i) demean-detrended, (ii) multiple regression to remove nuisance signals (i.e., white matter, CSF, mean-BOLD signal, detrended head realignment parameters, and the first-order derivatives of these variables), (iii) motion ‘scrubbing’^62^, where motion-contaminated volumes were flagged by framewise displacement (FD) greater or equal 0.2mm and replaced by interpolated volumes, (iv) bandpass filtering (0.009-0.08Hz), and (v) removal of interpolated frames that was only used to preserve time series order during bandpass filtering.

The preprocessed cortical functional data were mapped to the 32k_fs_LR surface using the single deformation map derived previously. The data on surface space is smoothed using a 6mm full-width-half-maximum (FWHM) Gaussian kernel. The subcortical functional data remained in volume space and was also smoothed with a 6mm FWHM Gaussian kernel. The cortical surface and subcortical volume data were combined into a CIFTI file^63^.

### 4.3. Parcellation and network construction

For each participant in MSC, an individualized parcellation and community assignments of functional networks were identified in ^7^. Boundary-mapping and watershed algorithm were used to identify the parcellation^13,57^, and the Infomap algorithm^64^ was used to estimate community assignments for each parcel (see ^7^ for detailed description).

A group-based parcellation atlas with corresponding functional network assignments is used in the present work: Gordon et al.^13^ (node *n* = 286; only nodes assigned to the major systems were included). Two additional parcellation schemes^14,22^ (Chan et al.^22^ and Schaefer et al.^14^) were used to verify whether the methods would work with group parcellations generated with other methods (see **SI**).

For each parcel, from either the individualized parcellation or the group-based parcellation, a parcel’s time series is computed as the average of the functional time series of all vertices within the parcel. For each of the *n*th individual, we created matrix denoted **H***_n_*, which is an *I_n_* individualized parcels *× J_n_* time points (TRs) matrix, and **G***_n_*, which is a *K* group parcels *× J_n_* time points (TRs) matrix, to represent the time series from individualized and group parcellation, respectively.

***MDS*** analyzes the correlation matrices computed from the time series data of the individualized parcellation (i.e., the **H***_n_* matrix) of each individual. ***CovSTATIS*** analyzes the correlation matrices computed from the time series of the group parcellation (i.e., the **G***_n_* matrices) of each individual. ***IndivSTATIS*** analyzes the cross-correlation matrices between the two types of time series (i.e., between **G***_n_* and **H***_n_*), resulting in a *K* group parcel *× I_n_* individualized parcel correlation matrix **X***_n_* for each of the *N* individuals. This matrix is called the *individualized connectivity block*. Unlike typical ‘functional correlation’ matrices, an individualized connectivity block **X***_n_* is a rectangular matrix that combines the information from an individual’s data extracted from the group parcellation and individualized parcellation. In all analyses, all correlation values of 1 are replaced with value of 0.99 to avoid introducing infinity values when performing Fisher’s *Z*-transformation (**SI** includes a version of the analysis with all negative correlations set to 0 and has other standard preprocessing steps when computing graph-based metrics, as detailed in ^22,65^.

### 4.4. IndivSTATIS and other multivariate methods

#### 4.4.1 Notation

We use bold uppercase letters to denote a matrix (e.g., **A**), bold lowercase letters to denote a vector (e.g., **a**), and italic letters to denote a scalar (e.g., *a* or *N*). Given an *N* (rows) *× M* (columns) matrix **X**, **x***_m_* is the *m*th column of the matrix, and *x_n,m_* is the value in the *n*th row and the *m*th column of **X**. A superscript ^T^ denotes the transpose of a matrix. **I** denotes the identity matrix, which has 1s on the diagonal and 0s off the diagonal. The operator trace() is the sum of all diagonal elements of a square matrix.

#### 4.4.2 IndivSTATIS

IndivSTATIS combines CovSTATIS and STATIS^26^ to jointly analyze all participants’ functional correlation patterns based on individualized parcellation, where each participant is represented by a *K* (group parcels) *× I_n_* (individualized parcels) correlation matrix **X***_n_*.

Each individualized connectivity block is normalized by its first eigenvalue (or called MFA-normalized^40^) to ensure equal contributions of all individuals to the extracted components.

IndiviSTATIS then computes the *K × K* cross-correlation matrix of each **X***_n_*, denoted **S***_n_*, where **S***_n_* = **X***_n_***X***_n_*^T^. From all **S***_n_*, IndivSTATIS quantifies the similarity between all **X***_n_* by computing the pairwise *R_V_* coefficients of all **S***_n_*. For example, the *R_V_* between **S***_n_* of the *n*th individual and **S***_n’_* of the *n’*th individual is computed as:

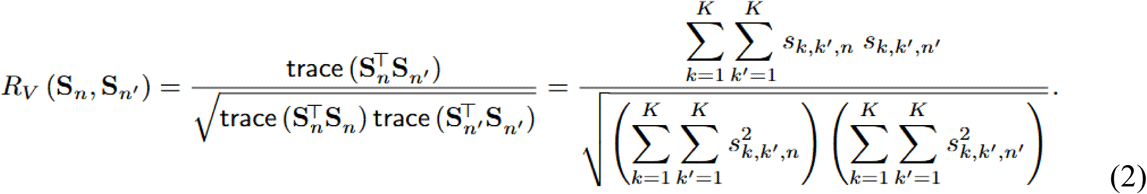

Note that *R_V_* coefficient is akin to a *squared* correlation coefficient and is, therefore, always positive. The set of *R_V_* coefficients are then stored in matrix **C**, where *c_n,n’_* is *R_V_* (**S***_n_*, **S***_n’_*). The first eigenvector of **C** (whose elements are always non-negative, see Perron-Frobenius theorem), denoted **u**_1_, is then extracted and normalized to have a sum of 1; the normalized weights are denoted *α*_n_ for each of the *n*th individualized connectivity block:

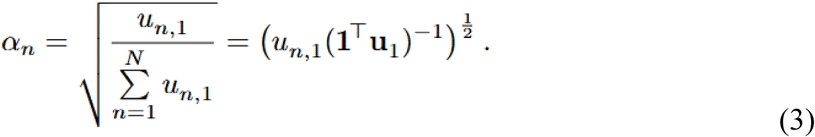

(note because the *α*_n_ are just a rescaling of **u**_1_, they are non-negative). The weighted blocks are then concatenated to create the *K* (group parcels) *× I* grand table, where 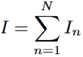:

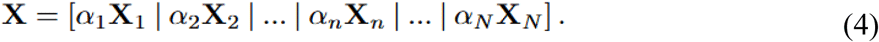

The grand table is then decomposed by the singular value decomposition (SVD):

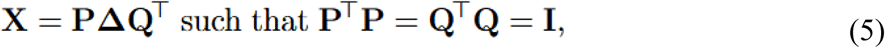

where **P** is an *K* (group parcels) *× L* (components) matrix of the left singular vectors, **Q** is the *I* (individualized parcels of all people) *× L* (components) matrix of the right singular vectors, and **Δ** is the *L × L* (components) diagonal matrix that stores the singular values *δ_l_* on the diagonal, where 1 ≤ *l* ≤ *L* and *δ*_1_ ≤ *δ*_2_ ≤ … ≤ *δ_l_* ≤… ≤ *δ_L_*. The grand table that entered the SVD included both positive and negative edges and the columns are not normalized. Negative values were included to preserve the difference between negative and zero connectivity. The columns are not centered (i.e., subtracting the mean), because zero brain connectivity is meaningful, and they are not scaled (i.e., divided by standard deviation or other variance measures), because these connectivity measures are on the same scale (i.e., -1 to 1) and their original variability includes important parcel-to-parcel relationships for building the component space.

The column factor scores that are associated with the *n*th participant **F**_[*n*]_ are then computed by

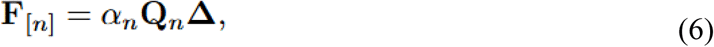

where **F**_[*n*]_ is *I_n_* (individualized parcels of the *n*th participant) *× L* (components) matrix, and the *l*th column of **F**_[*n*]_ (denoted by **f**_[*n*]*l*_) represents all individualized parcels on the *l* th component.

#### 4.4.3 MDS and CovSTATIS

To validate IndivSTATIS results, we compared the resulting components with two unsupervised multivariate methods: MDS and CovSTATIS. MDS is usually applied to distance matrices by transforming a distance matrix to a positive semi-definite matrix. However, as connectivity matrices are already positive semi-definite, MDS in this context simply performs an eigenvalue decomposition (EVD) on each single non-normalized (i.e., not centered or scaled) correlation matrix (**X***_n_*) of the *n*th participant. We did not normalize the connectivity matrix because 1) a correlation value of 0 is meaningful (i.e., not centering will preserve the origin) and 2) all connectivity measures share the same scale (i.e., correlation coefficients range between -1 and 1). MDS extracts a component space for each **X***_n_*, with factor scores representing the position of each brain parcel in the component space. The components from MDS are ordered by the amount of variance they explain, in a descending order.

CovSTATIS performs a joint component-based analysis across multiple connectivity matrices with the same matrix dimensions. Specifically, CovSTATIS first normalized each connectivity matrix by their first eigenvalue to ensure equal contributions from all participants when extracting dimensions. Next, CovSTATIS computes the structural similarity, measured by *R_V_* coefficients (equation 2), between all connectivity matrices and produces an *n × n* matrix **C**. CovSTATIS derives weights for each connectivity matrix from the scaled (to sum to 1) first eigenvector of **C** (equation 3). Matrices exhibiting more common patterns are weighted more, whereas matrices with rarer patterns are weighted less, limiting the impact of outlier matrices. CovSTATIS applies these weights to compute the optimal linear combination—a weighted average called the *compromise* matrix:

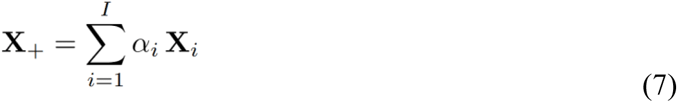

and extracts component with an EVD:

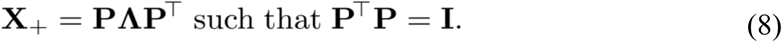

The factor scores (**F**) represent the brain configuration across all participants (i.e., group-level brain configuration) and are computed as:

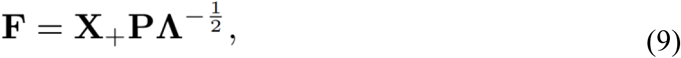

and the connectivity pattern of each participant can be projected onto the same space as *partial factor scores*, extracting individual-level brain configurations:

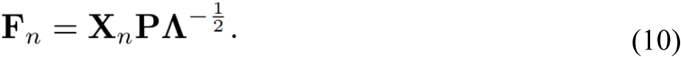

Together, the weighted average of these partial factor scores (representing the individual-level brain configurations) gives the factor scores (representing the group-level brain configuration):

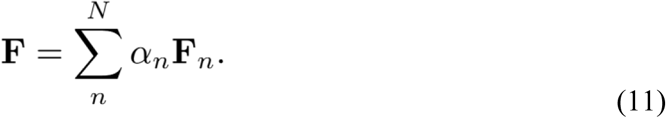

## 4.5. Data/code availability statement

All analysis code used in this study is available via the Open Science Framework (OSF) at: https://osf.io/ybwt9/ and https://github.com/juchiyu/riip. The neuroimaging data were obtained from the Midnight Scanning Club (MSC) dataset, which is publicly accessible via the MSC GitHub repository: https://openneuro.org/datasets/ds000224/versions/1.0.4

## 4.6. Ethics statement

This study used the publicly available Midnight Scan Club (MSC) dataset (OpenNeuro ds000224). The original data collection procedures were approved by the Institutional Review Board (IRB) of Washington University in St. Louis. All participants provided informed consent, including consent for their de-identified data to be shared publicly for research purposes. With secondary analysis of the MSC data, this study does not require new IRB approval.

## 4.7. Disclosure of competing interests or affirmative statement

The authors declare no competing interests.

## Acknowledgements

J-CY receives grant support from the Womenmind postdoctoral research award of the Centre for Addiction and Mental Health (CAMH). EWD has received funding from BBRF, NIMH, CIHR, and CAMH Foundation. The authors have no other relevant financial relationships or other conflicts of interest to disclose.

