## Supplementary material for "IndivSTATIS: A multivariate approach to analyze brain network configurations with individualized parcellation": SI

### 1. Supplemental Results

#### 1.1 IndivSTATIS results with no negative edges

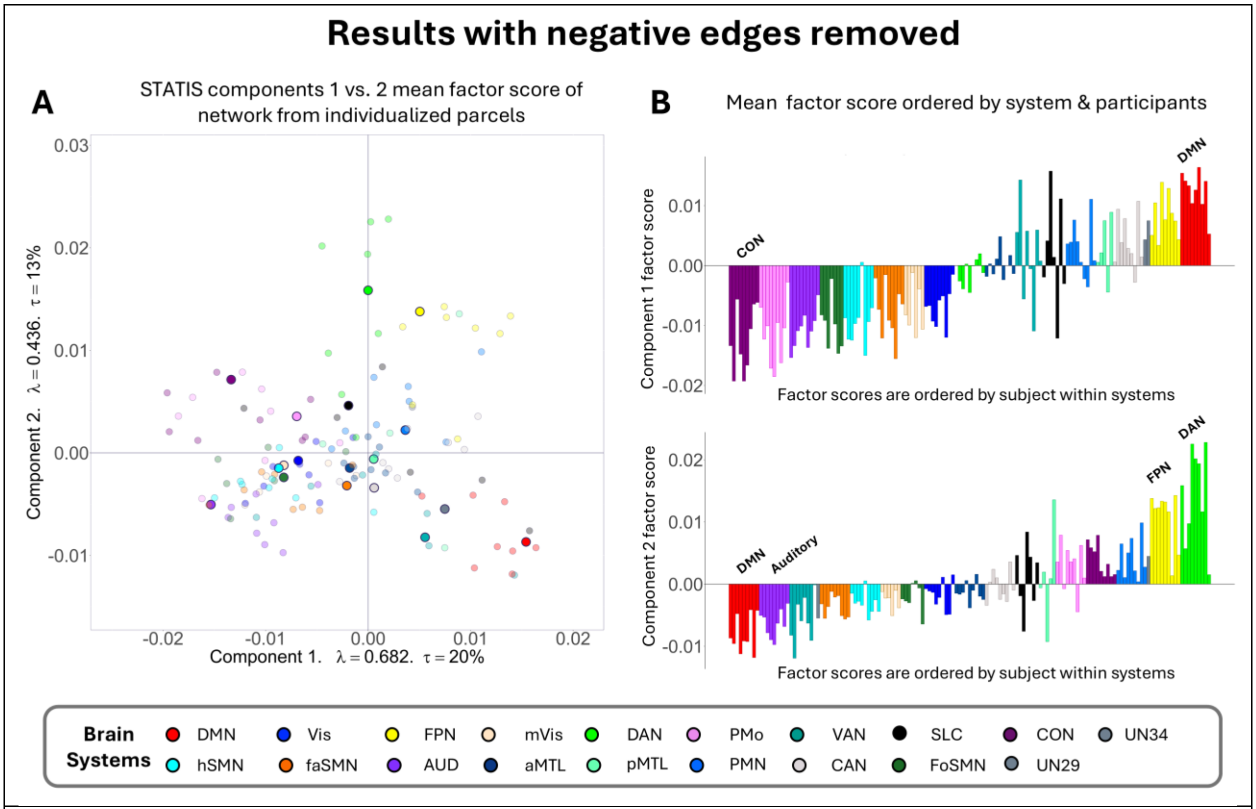


**Figure S1 – IndivSTATIS excluding negative edges.**
Similar to Figure 3 in the main manuscript. **(A)** Participants' mean column factor score for each functional network are plotted along IndivSTATIS Components 1 and 2. To single out one participant, MSC-01's functional network's mean column factor scores are plotted as large and solid circles; all the other participants are plotted as smaller and transparent circles. Component 1 is characterized by the opposition, indicating the anticorrelation, of two association networks: cingulo-opercular (CON; dark purple) and default mode network (DMN; red; see main manuscript Figure 1 for topographic location of CON and DMN across participants). **(B)** The barplots (top: Component 1; bottom: Component 2) sort each participant's mean factor score by (i) network and (ii) participant number. In Component 1, CON [purple] is most negative across all participants, so it is grouped together on the left-most side of the barplot, and MSC-01's CON mean factor score is the first purple bar.

### 1.2 Comparisons between IndivSTATIS, MDS, and CovSTATIS with all 9 participants

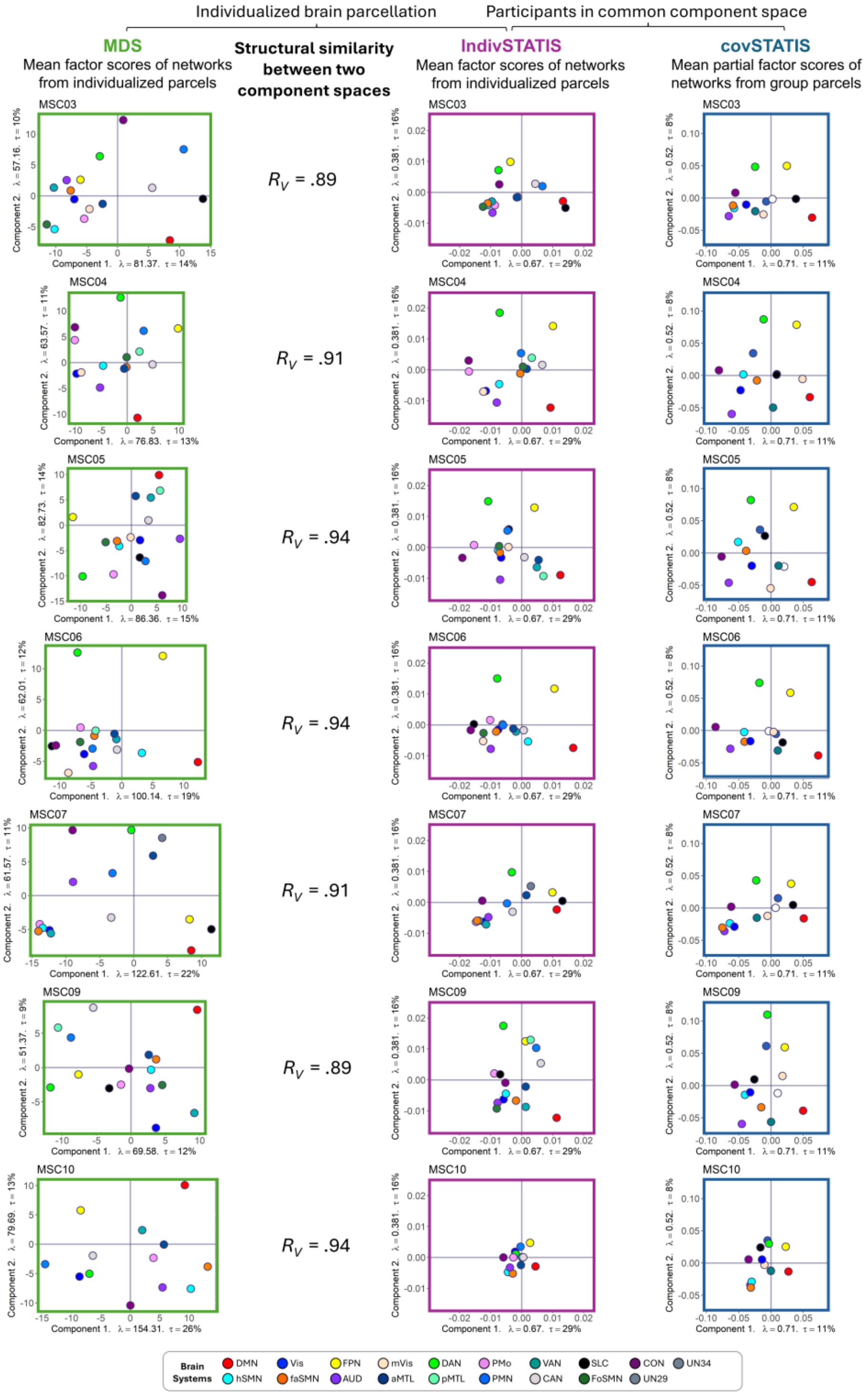


**Figure S2 – Comparisons to MDS and covSTATIS for all MSC participants.**
Component biplots for MSC03 to MSC10 (excluding MSC08 for poor quality data) are presented to illustrate the differences in network configuration between these methods. This figure shows the results analyzing the data derived from the same fMRI scan using

multidimensional scaling (MDS; green boxes in the left column) and IndivSTATIS (purple boxes in the middle column) with individualized networks, and covSTATIS (blue boxes in the right column) with networks derived from a group atlas (i.e., Gordon et al. 2016). Each round dot illustrates the mean factor score of each network and is colored accordingly. The $R_V$ coefficients, an $R^2$ analogous measure ranging from 0 to 1, quantify the structural similarity between component spaces derived from MDS and IndivSTATIS, both based on individualized parcellation. The results showed that both the individualized spaces from MDS and the participant alignment from covSTATIS were well preserved when analyzed jointly by IndivSTATIS.

#### 1.3 IndivSTATIS using different common references for parcels

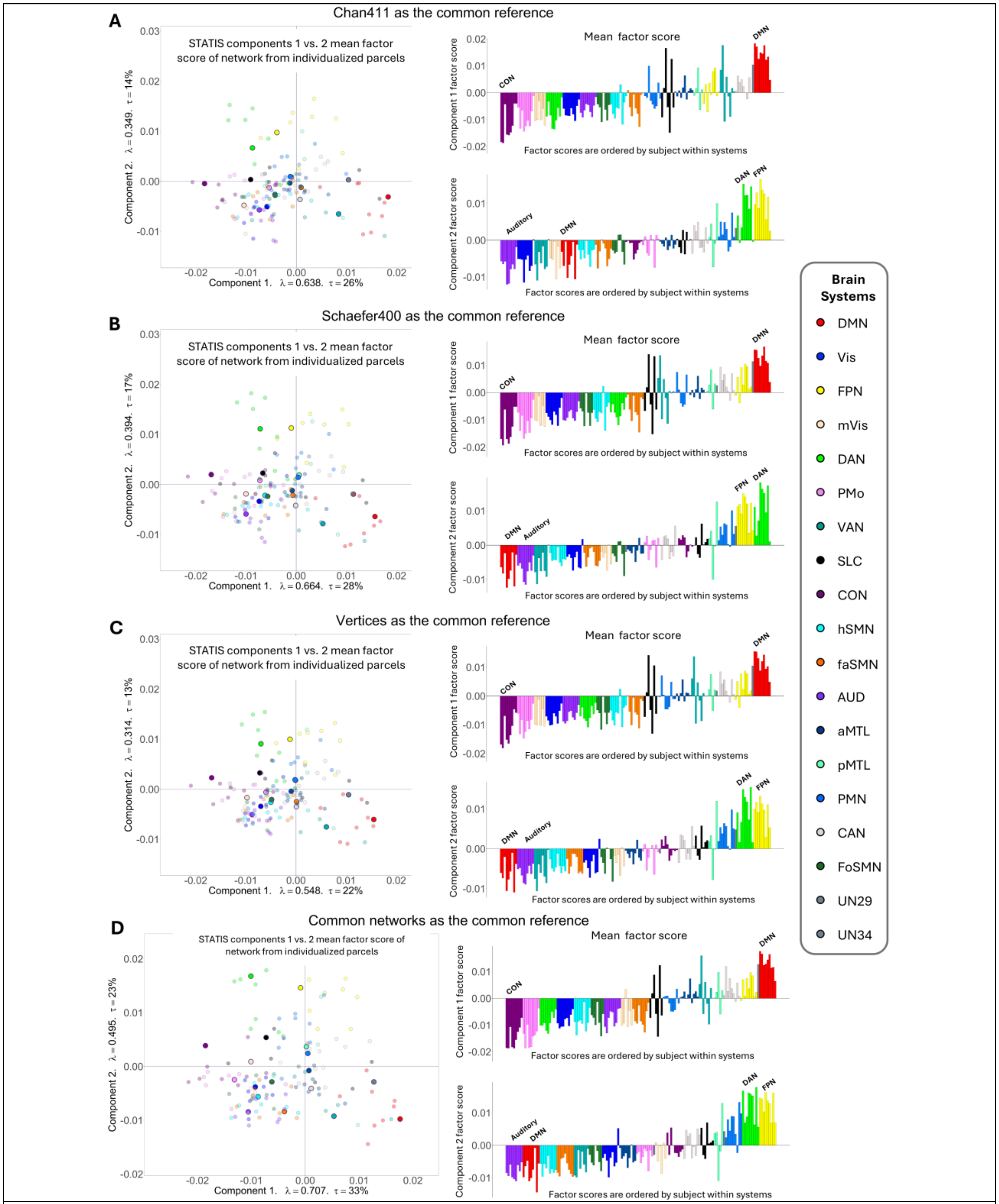


**Figure S3 – IndivSTATIS results with different group references.**
Column factor scores from IndivSTATIS using four different types of group references: (**A**) Chan411 group parcellation, (**B**) Schaefer 400 group parcellation, (**C**) vertices, and (**D**) the

common networks across all 9 participants. The results showed similar factor structure across different approaches with the current data. In each plot, the left panel showed the first two IndivSTATIS column factor scores on the first two components. To highlight one participant, MSC-01's functional network's mean column factor scores are plotted as large and solid circles; all the other participants are plotted as smaller and transparent circles. All dots are colored according to their networks. The right panel of each plot showed barplots (top: Component 1; bottom: Component 2) where each participant's mean factor scores were sorted by (i) network and (ii) participant number. In other words, on Component 1, cingulo-opercular network (CON; purple) [purple] is most negative and default mode network (DMN; red) is the most positive across all participants, so it is grouped together on the left-most side of the barplot. MSC-01's CON mean factor score is the first purple bar, and MSC-10's CON mean factor scores is the last purple bar.
